# Distinct Proline residues in hinge-regions of SYMRK generates a phosphocode for releasing Malectin-like-Domain to allow progress of rhizobia-legume symbiosis at epidermal-cortical barrier

**DOI:** 10.1101/2024.08.14.607956

**Authors:** Dipanjan Chakrabarti, Anindita Paul, S. Bhattacharyya, Sagnik Das, Firoz Molla, Alokmoy Biswas, Maitrayee DasGupta

**Affiliations:** Department of Biochemistry, University of Calcutta, 35, B. C. Road, Kolkata 700019, India

**Author notes:** D.C. and A.P. contributed equally to this work. One Sentence title: Signature prolines that distinguishes SYMRK from MLD-LRR-RLKs evolved a phosphocode for releasing MLD to allow symbionts through transcellular barriers during intracellular rhizobia-legume symbiosis. The author(s) responsible for distribution of materials integral to the findings presented in this article in accordance with the policy described in the Instructions for Authors (https://academic.oup.com/plcell/pages/General-Instructions) is: Maitrayee DasGupta.

## Abstract

Symbiosis Receptor Kinase (SYMRK), a malectin-like-domain/leucine-rich-repeat receptor-like-kinase (MLD-LRR-RLK), is the upstream most component in Common-Symbiosis-Signalling-Pathway. We highlight two Proline residues that were distinctly acquired by SYMRK orthologues in its hinge-regions to evolve a signalling module for allowing progress of symbionts across transcellular barriers during rhizobia-legume symbiosis. Within the Ectodomain hinge (EctoD-hinge) all MLD-LRR-RLKs have a conserved W^1^x_n_GDPCx_n_W^2^x_4_C motif, where SYMRK orthologues within legumes have a distinct signature with a Proline preceding W^2^ enabling cleavage of SYMRK-EctoD for releasing MLD. Within the kinase hinge (KD-hinge) at gatekeeper+1 position, a conserved Glutamate in MLD-LRR-RLKs is replaced by Proline in all SYMRK orthologues that enabled its dual-specific kinase activity for ensuring EctoD-cleavage. Substitution of either Proline restricted symbionts at the transcellular epidermal-cortical barrier forming infection patches in the nodule apex without affecting epidermal invasion and nodule organogenesis. This halt was entirely overcome by ectopic expression of free MLD demonstrating the released MLD to have an active role in progress of symbionts. Overall, we show that adaptations of distinct Prolines in hinge-regions of SYMRK orthologues in legumes have evolved a signalling module involving dimerization and optimal phosphorylation of SYMRK for releasing MLD as an active transducer of symbiosis signalling.

## Introduction

CSSP co-evolved with intracellular symbiosis of plants with phosphate-acquiring Glomeromycotean fungi when plants colonised land 450 Mya. (Delaux et al., 2015; Parniske, 2008). More recently in evolution by 92–110 Mya, CSSP was co-opted for intracellular RNS with nitrogen fixing bacteria within plants belonging to a monophyletic NFC comprising of four orders Fabales, Fagales, Cucurbitales and Rosales (Doyle, 2011). Members of two classes of receptors are central to RNS signalling: the LysM-RLKs, (NFR1/NFR5, LYK3/NFP) (Madsen et al., 2003; Smit et al., 2007) and a MLD-LRR-RLK, (SYMRK/DMI2) (Endre et al., 2002; Stracke et al., 2002). Activation of SYMRK following perception of symbionts by LysM receptors, leads to perinuclear calcium oscillations (Kosuta et al., 2008). The calcium signature is decoded by CSSP members: calcium/calmodulin-dependent kinase (CCaMK/DMI3) (Lévy et al., 2004) and a transcriptional regulator CYCLOPS/IPD3 (Messinese et al., 2007; Yano et al., 2008) for activating RNS specific master transcriptional regulator NODULE INCEPTION (NIN) to initiate symbiosis (Singh et al., 2014). Trans-complementation of the angiosperm mutants with orthologs of CSSP members from diverse species indicated functional conservation of CSSP for supporting intracellular mutualistic symbioses across the embryophytes (Das et al., 2019; Delaux et al., 2015; Gherbi et al., 2008; Godfroy et al., 2006; Markmann et al., 2008; Saha et al., 2014; Yano et al., 2008).

SYMRK is the upstream-most member of CSSP and is described as an enigmatic receptor that guard and guide microbial endosymbioses with plant roots (Holsters, 2008; Limpens et al., 2005). Symbiont specificity and compatibility in RNS do not depend on the origin of SYMRK (Gherbi et al., 2008; Markmann et al., 2008; Saha et al., 2014), rather it is believed to thwart off the mechanical stress or touch responses in host plant in presence of symbionts (Esseling et al., 2004). SYMRK along with its interactors/substrates are indispensable for the progress of symbiosis at both epidermal and cortical levels and plays an important role in synchronizing the responses at both levels (Chen et al., 2012; Den Herder et al., 2012; Kevei et al., 2007). At the epidermis rhizobia gets entrapped by a root hair curl and then are guided by formation of infection threads (IT) that transcellularly migrates toward a nodule primordium in the cortex (Fournier et al., 2015). In the cortex the endocytic uptake of symbionts into organelle-like structures known as symbiosomes is thought to be the key step during the evolution of rhizobia-legume symbiosis (Parniske, 2018).

SYMRK has an active intracellular kinase-domain (KD) and loss of kinase activity leads to loss of all symbiotic features (Yoshida & Parniske, 2005). Like several other RLKs, SYMRK gets phosphorylated in the gatekeeper tyrosine residue in the hinge-region in the core-KD (Oh et al., 2009; Samaddar et al., 2013; Suzuki et al., 2016), and this phosphorylation was important for synchronising the epidermal and cortical symbiotic program (Saha et al., 2016). Phosphorylation outside the core-KD in the C terminal end is important for nodule organogenesis (Abel et al., 2024), whereas SYMRK core-KD devoid of intramolecular autoinhibitory interactions trigger spontaneous hypernodulation (Saha et al., 2014). Overexpression of full-length SYMRK can also prompt spontaneous nodule organogenesis and such organogenesis triggered through forced interaction between NFR1/NFR5 relies on SYMRK (Ried et al., 2014; Rübsam et al., 2023). A constitutive cleavage of SYMRK ectodomain was noted at a conserved GDPC containing motif between MLD and LRRs and the *Lotus japonicus* symRK-14 with a mutation in GDPC motif uncouples the epidermal and cortical symbiotic program (Antolín-Llovera, Ried, et al., 2014; Kosuta et al., 2011). Since this cleavage site of SYMRK is positioned within and not at the stalk of the ectodomain it is distinct from ectodomain shedding described for receptor tyrosine kinases. The ectodomain cleavage of SYMRK is restricted in presence of rhizobia and stabilisation of full length receptors is a key step for rhizobial invasion (Pan et al., 2018). Mutations in the conserved residues within MLD makes SYMRK susceptible to proteasomal degradation and cannot restore symbiosis in *symrk null* mutants (Pan et al., 2018).

How ancient CSSP started functioning or how different forms of symbiosis evolved by co-opting CSSP is a fascinating puzzle that is yet to be deciphered. In this article, we aim to highlight the molecular adaptations in SYMRK, that have contributed towards evolving rhizobia-legume symbiosis. We used biochemical and cell biological approaches to decipher the functional significance of two distinct Proline residues adapted in hinge-regions of SYMRK orthologues. It is apparent that specific Proline adaptations in SYMRK led to evolution of a signalling module where dimerization and optimal phosphorylation of its Kinase domain drives ectodomain cleavage and release of MLD to allow transcellular migration during rhizobia-legume symbiosis.

## Results

### SYMRK orthologues have distinct Proline residues adapted in hinge-regions

SYMRK orthologues form a distinct clade within MLD-LRR-RLKs representing LRR-RLK-I, and the clustering within SYMRK was consistent with species tree highlighting the NFC (Fig. 1, A and B, and Supplementary Fig. S1A). Two distinct Proline residues within hinge-regions distinguished SYMRK orthologues from MLD-LRR-RLKs which is in accordance with hinge motions playing a significant role in the functional evolution of proteins (Campitelli et al., 2020). The intrinsically flexible KD-hinge connecting N and C-lobe is required for Kinase activation (Chen et al., 2007), and almost all RLKs have a Tyr residue in gatekeeper position adjacent to KD-hinge (Shiu & Bleecker, 2001). Various evidence indicate that protein kinases are regulated by different mechanisms controlled primarily by their respective gatekeeper residues (Azam et al., 2008; Chen et al., 2007; Emrick et al., 2006). SYMRK orthologues within or outside NFC distinctly have a Proline in the gatekeeper+1 position where the rest of the MLD-LRR-RLKs have Glutamate (Fig. 1C and Supplementary Fig. S2, A and B). Even SYMRK orthologues from *Marchantia paleacea*, *Selaginella moellendorffii* have this Proline conserved (Supplementary Figs. S1A and S2A), indicating its importance since the adaptation of SYMRK within CSSP for mycorrhizal symbiosis (Radhakrishnan et al., 2020). There was one MLD-LRR-RLK (PTQ39574.1) with a Proline in gatekeeper+1 position, from *Marchantia polymorpha* a nonsymbiotic liverwort that does not have a SYMRK orthologue. In a sequence similarity tree developed separately with KD but not with EctoD, this kinase PTQ39574.1 clustered within SYMRK orthologues (Supplementary Fig. S1, B and C). Zygnematophyceae, the algal sisters of land plants have proper orthologues for CCaMK/CYCLOPS but for SYMRK there is a pro-orthologue (Zci_05951) with Glutamine in gatekeeper+1 position (Feng et al., 2024). Arabidopsis MLD-LRR-RLK (NP_564904.1) that failed to complement *symrk null* mutant had a Glutamate in gatekeeper+1 position (Li et al., 2018). It appears that adaptation of Proline in gatekeeper+1 position within SYMRK have contributed towards driving the evolution of CSSP in land plants. SYMRK resembles LRR-RLK-II members after cleavage of its ectodomain and other than SYMRK, Proline in gatekeeper+1 position is predominant only in LRR-RLK-II, suggesting an overlap in their origin/function (Supplementary Fig. S2C). But overall, Glutamate is the most prevalent residue in Gatekeeper+1 in all LRR-RLKs and a strong preference for Glutamate is also noted within human kinome for stabilizing the hinge (Brooijmans et al., 2010). Since backbone flexibility within the KD-hinge is essential for kinase activation, the conformational rigidity that a Proline brings to the hinge-region of SYMRK-KD, appeared necessary for its functional adaptation.

**Figure 1.**
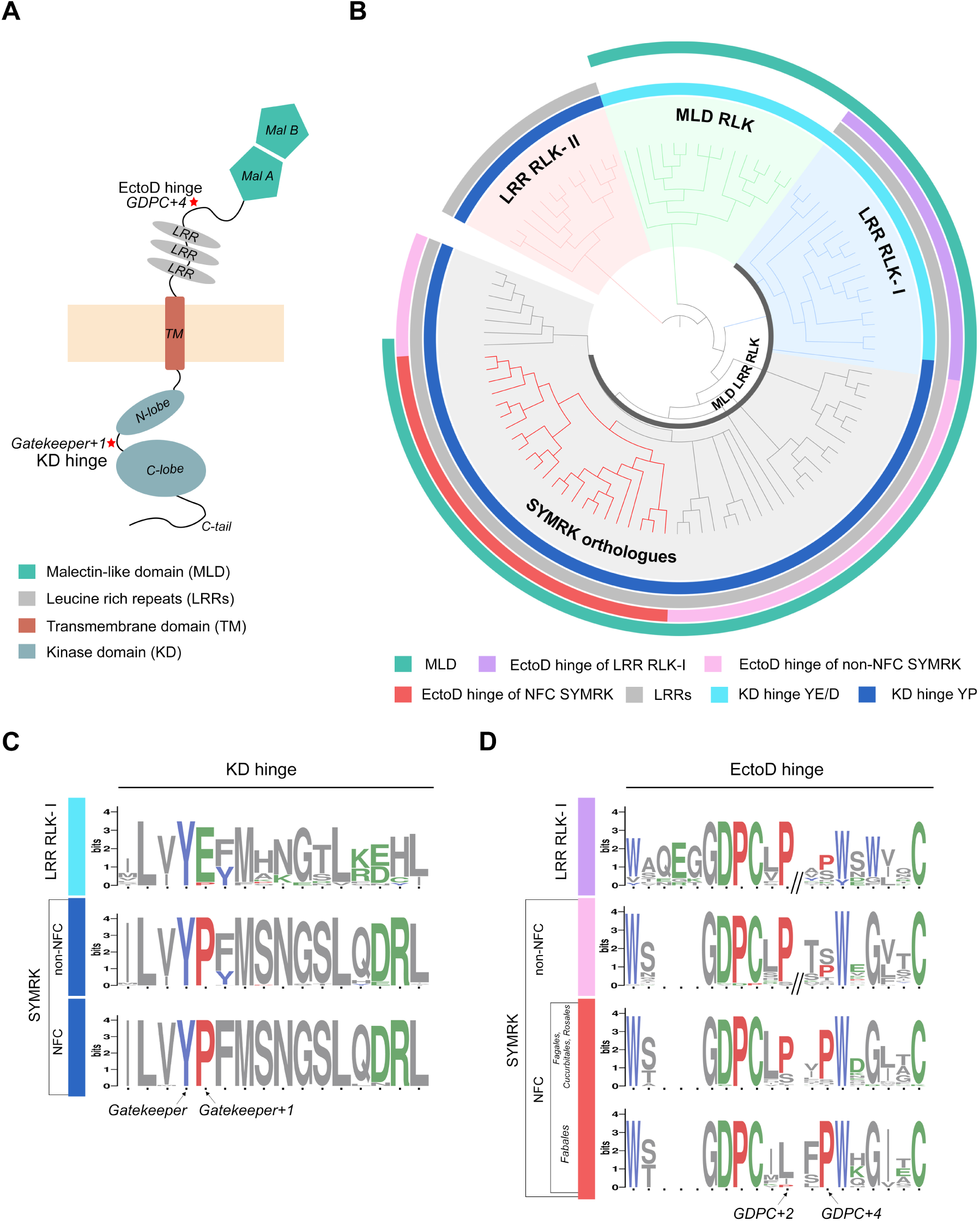
SYMRK orthologues have conserved Proline residues both in KD hinge and EctoD hinge. (**A**) Domain structure of SYMRK. Asterisks indicate the hinge regions of the kinase domain (KD-hinge) and the ectodomain (EctoD-hinge). (**B**) Maximum likelihood sequence similarity tree of representative members from MLD-RLK, LRR-RLK-II and MLD-LRR-RLKs including SYMRK orthologues of LRR-RLK-I across embryophytes. Clades are color labeled as indicated and NFC is highlighted in ‘red’. The outer 3 color rings indicate, LRR, variation in EctoD-hinge and MLD. The innermost ring indicates the variation in KD-hinge. For detail, see Supplementary Fig.S1. (**C** and **D**) Sequence logos demonstrate divergence of KD-hinge and EctoD-hinge in SYMRK orthologues (NFC and non-NFC) and other MLD-LRR-RLKs from LRR-RLK-I. Gatekeeper, gatekeeper+1 positions in KD-hinge (C) and GDPC+2, GDPC+4 positions in EctoD-hinge (D) are indicated. For alignments, see Supplementary Figs. S2 and S4.

In the EctoD of SYMRK, the linker that connects MLD and LRRs is predicted to be a potential hinge-region which contains a motif of W^1^x_n_GDPC^1^x_n_W^2^x_4_C^2^ shared by most MLD-LRR-RLKs (Supplementary Fig. S3). AlphaFold2 derived static structures of several MLD-LRR-RLKs including SYMRKs indicated the structural conservation of this motif (Supplementary Fig. S4C). However, the nature and length of insertions within the conserved residues varied between MLD-LRR-RLKs, where SYMRK orthologues within NFCs have a signature of W^1^xGDPC^1^x_3_-_4_W^2^xGx_2_C^2^ (Fig. 1D and Supplementary Fig. S4). In GDPC+2 position Proline was predominant in most MLD-LRR-RLKs, as well as in SYMRKs from non-NFC and in some non-legumes within NFC. But a Proline just preceding W^2^ in GDPC+4 position was a distinctive feature of SYMRK from NFC and was seldom noted in non-NFC. In Fabales, with the exception of caesalpinoids and mimosoids, there was a conspicuous absence of Proline in the GDPC+2 position giving a distinct EctoD-hinge signature for the SYMRK orthologues in papilionoids. It may be noted that SYMRKL1 (LotjaGi2g1v0191100), a GPI anchored protein from *Lotus japonicus* (Frank et al., 2023), as well as the pro-orthologue of SYMRK in Zygnematophyceae (Zci_05951) (Feng et al., 2024), do not have the consensus noted within SYMRK orthologues. Taken together, a Proline preceding W^2^ in GDPC+4 position is acquired in SYMRK orthologues of Fabales where RNS evolved as a stable genetic trait in rhizobia-legume symbiosis. The fact that the Proline within GDPC motif was essential for EctoD cleavage in SYMRK (Antolín-Llovera, Ried, et al., 2014), we reasoned that the adapted Proline at GDPC+4 could be involved in the same process. The overall robust and distinctive adaptation of Prolines in KD-hinge and Ecto-hinge of SYMRK orthologues prompted us to investigate whether these changes in hinge regions were related to functional adaptation of SYMRK during rhizobia-legume symbiosis.

### Gatekeeper+1 Proline in KD-hinge required for optimal activation of SYMRK and crossover of symbionts at the epidermal-cortical barrier

Earlier, we had established the significance of gatekeeper tyrosine phosphorylation in the KD-hinge within the core-KD using SYMRK from *Arachis hypogaea* (hereafter SYMRK) (Saha et al., 2016; Samaddar et al., 2013). SYMRK-KD is a dual-specific Ser/Thr/Tyr kinase that autophosphorylated in gatekeeper Y670 (residue numbering from *Arachis* SYMRK) and generated a phosphocode of pS754↑pS757↓pT763↑pY766↑ in its activation loop (Bhattacharya et al., 2019). In contrast, Y670F-SYMRK-KD with a nonphosphorylatable gatekeeper was basally phosphorylated and the phosphorylation was restricted to Ser/Thr residues with a distinct phosphocode of pS754↓pS757↑pT763↓pY766↓. To understand the significance of conservation of gatekeeper+1 Proline we substituted it with Glutamate (P671E), the most prevalent residue in this position in all LRR-RLKs (Supplementary Fig. S2C). Unlike WT-SYMRK-KD that autophosphorylated in Ser/Thr/Tyr, both P671E-SYMRK-KD and Y670F-SYMRK-KD were active kinase and autophosphorylated only on Ser/Thr (Fig. 2A). Gatekeeper Tyr (Y670) phosphorylation was abolished in P671E-SYMRK-KD and pTyr was undetectable in both the KD-hinge mutants. Also unlike optimal phosphorylation in WT-SYMRK-KD both KD-hinge mutants were basally phosphorylated (Fig. 2B). LC-MS/MS derived abundance of phosphopeptides revealed the autoactivation sites of P671E-SYMRK-KD to resemble that of Y670F-SYMRK-KD generating a phosphocode that is distinct from WT-SYMRK-KD (Fig. 2C and Supplementary Data Set 1). Thus, the conserved P671 in gatekeeper+1 position determines phosphorylation in gatekeeper Y670 and the dual specificity of kinase activity of SYMRK-KD to generate a distinct phosphocode in core-KD.

**Figure 2.**
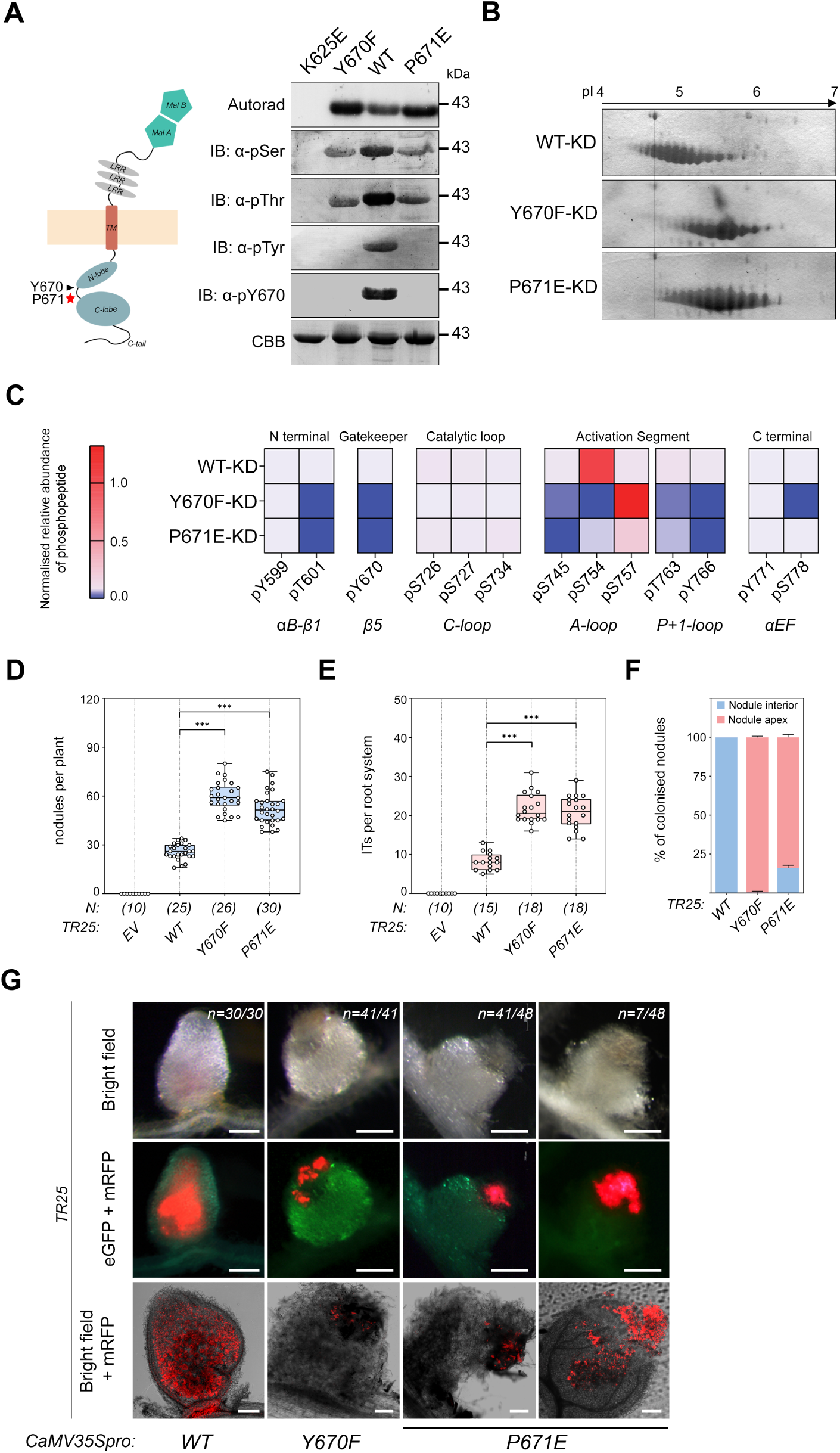
Significance of gatekeeper+1 Proline in SYMRK KD-hinge. (**A-C**) Comparing autoactivation of gatekeeper+1(P671E-KD) with gatekeeper (Y670F-KD) mutants of 6xHis tagged SYMRK. (**A**) *In vitro* autophosphorylation (^32^P autorad) and western blots using α-pSer, α-pThr, α-pTyr, α-pY670 antibodies of KD-hinge mutants. Similar to Y670F-KD, Tyrosine phosphorylations were abolished in P671E-KD. Protein abundance visualized by Coomassie Brilliant Blue staining (CBB). The catalytically inactive K625E-KD is used as reference. (**B**) KD-hinge mutants were analyzed by isoelectric focusing on Immobiline DryStrips, pH 4 to 7 followed by SDS-PAGE and visualized by staining with Coomassie Brilliant Blue G 250. A reference point for all three 2D gels is indicated by the dotted line. P671E-KD and Y670F-KD polypeptides had a basal level of phosphorylation (pI ∼5.5) as compare to optimally phosphorylated WT-KD (pI ∼5). (**C**) Heatmap shows the normalized LC-MS/MS derived relative abundance of phosphopeptides indicating the phoshosites in KD-hinge mutants. The phosphocode generated in the WT-KD differs from the phosphocode generated in P671E-KD and Y670F-KD. The smallest and largest means within each dataset are identified as 0.0 and 1.0 respectively. This heatmap is a representative of two independent experiments (see Supplementary Data Set 1). (**D-G**) Effect of Y670F and P671E mutation on nodule development and symbiont colonization. *TR25* complemented with *WT-SYMRK-eGFP*, *Y670F-SYMRK-eGFP* and *P671E-SYMRK-eGFP* and infected with *Sinorhizobium meliloti.* IT and Nodule development were monitored in transgenic hairy roots at 1WAI and 6WAI respectively. Box whisker plots represent the total number of nodules/ plant (**D**) and the number of ITs/ root system (**E**). An average of 3 biological replicates each having n > 8 plants for nodule count and n > 4 plants for IT count were analyzed. The number of plants (N) analyzed for each construct are indicated beneath the respective box. Error bar denotes standard deviation (SD). Asterisks denote statistical significance as calculated by using an unpaired two-tailed t test: ***p < 0.001. (**F**) Histogram represents the distribution of rhizobial colonization in nodule apex and nodule interior. (**G**) Stereo images shown as bright-field (upper panels), merged images of eGFP and mRFP (middle panels) and confocal images of merged mRFP and bright field (lower panel). The letter ‘n’ indicates the total number of nodules analyzed for each case. Scale bars, for upper and middle panel, 500μm; for lower panel, 40μm. For details, see Supplementary Fig. S5.

SYMRKs are not involved in determining rhizobial preferences and cross-species complementation can restore symbiosis in *symrk-null* mutants of model legumes (Gherbi et al., 2008; Markmann et al., 2008). We have reported successful complementation of *Medicago symrk−/− TR25* by expression of *Arachis hypogaea* SYMRK from constitutive CaMV35S promoter (Saha et al., 2014; Saha et al., 2016). To assess the functional significance of distinct adaptations of Proline residues in SYMRK, we continued heterologous complementation in *TR25* under identical conditions using CaMV35S promoter for the entire investigation. Ectopic expression would highlight the checkpoints that are strictly guarded by kinase activity at post translational level and avoid possible defects in symbiosis that can be displayed by suboptimal protein levels while expressing SYMRK mutants with the native promoter (Antolín-Llovera, Ried, et al., 2014). The nodule counts as well as the IT counts in P671E-SYMRK:*TR25* was significantly higher than WT-SYMRK:*TR25* (Fig. 2, D and E, and Supplementary Table S1). But the distinguishing feature in P671E-SYMRK:*TR25* was the arrest of IT progression at the epidermal-cortical interface, resulting in extensive infection patches at the nodule apex (n= 41/48) (Fig. 2, F and G, and Supplementary Fig. S5C). Also, the ITs had inflated sac-like structures unlike WT-SYMRK:*TR25* where there was successful complementation and normal IT progression (Supplementary Fig. S5, A and C). In a few cases of P671E-SYMRK:*TR25* (n=7/48), ITs slightly ramified inside the nodule interior but there was no release of rhizobia, and the nodules were always white. These observations resemble Y670F-SYMRK:*TR25*, where also rhizobial progression was stalled in the epidermal-cortical barrier and like P671E-SYMRK:*TR25* the nodule counts as well as the IT counts were significantly higher than WT-SYMRK:*TR25* (Fig. 2, D and E, and Supplementary Fig. S5, B and C, and Supplementary Table S1). These high counts reflected the host plant’s attempt to compensate for anomalies in IT propagation and nodule development (Kuppusamy et al., 2004). Taken together, the distinct gatekeeper+1 Proline determines the requisite activation state of KD for allowing rhizobial progression beyond the epidermal-cortical barrier. This also indicates that the basal phosphocode in SYMRK was sufficient for the initiation of mutualistic interaction between rhizobia and legumes at the epidermis that led to IT development and nodule organogenesis. Thus functional involvement of SYMRK at the epidermal-cortical barrier is distinct from its involvement in the epidermis.

### GDPC+4 Proline in EctoD-hinge required for Ectodomain cleavage of SYMRK and crossover of symbionts at the epidermal-cortical barrier

SYMRK undergoes constitutive cleavage within the conserved GDPC motif of EctoD-hinge but stabilization of full-length receptors was essential for rhizobial invasion (Antolín-Llovera, Ried, et al., 2014; Pan et al., 2018). To understand the significance of a conserved Proline in the GDPC+4 position, we developed an *in vitro* cleavage assay with SYMRK-EctoD expressed in *E.coli* and the apoplastic sap of *Arachis hypogaea* roots. Since there is an onset of a major transcriptional program at 4 days post infection (4dpi) during the primordia formation in *Arachis hypogaea* roots (Karmakar et al., 2019), we have used apoplastic extracts from uninfected (0dpi) and infected (4dpi) roots as a source of proteases. The EctoD cleavage was monitored using anti-MLD (Fig. 3, A and B) and confirmed by LC-MS/MS (Supplementary Fig. S6, A and B and Supplementary Data Set 2). A mild cleavage releasing MLD was noted without adding any apoplast which could be intrinsic autocatalytic cleavage of SYMRK-EctoD as mentioned earlier (Antolín-Llovera, Ried, et al., 2014). Apoplast treatments further activated the cleavage where the 4dpi-apo was significantly more efficient (Fig. 3B). This indicates SYMRK-EctoD cleavage to be an enzymatically driven active process that is important during the progress of symbiosis. The cleavage was then tested with P393L-SYMRK-EctoD with substitution in GDPC+4, along with P388L-SYMRK-EctoD with substitution in GDPC motif, and a double mutant P388L/P393L-SYMRK-EctoD (Fig. 3, C and D). As compared to WT-SYMRK-EctoD, the cleavage of all three mutant SYMRK-EctoDs was affected, with the efficacy of cleavage being WT>P388L>P393L >P388L/P393L. We then attempted to observe the cleavage in *Nicotiana benthamiana* leaves. This was demonstrated before with *Lj*SYMRK and was thought to be mediated by ubiquitously present proteases or it could be an intrinsic auto-cleavage activity of the ectodomain (Antolín-Llovera, Ried, et al., 2014). In contrast to WT-SYMRK, where the full-length receptor was not detectable in *Nicotiana* leaves, in both the KD-hinge (Y670F, P671E) and EctoD-hinge (P388L, P393L) mutants we could detect the intact receptor, indicating EctoD cleavage to be restricted in hinge mutants (Supplementary Fig. S6B). The restriction in cleavage in EctoD-hinge mutants confirmed the importance of P393 (GDPC+4) along with P388 (GDPC) in determining the cleavability of SYMRK ectodomain. But most importantly the restriction of cleavage in KD-hinge mutants indicated the activation state of SYMRK-KD to have a role in EctoD cleavage.

**Figure 3.**
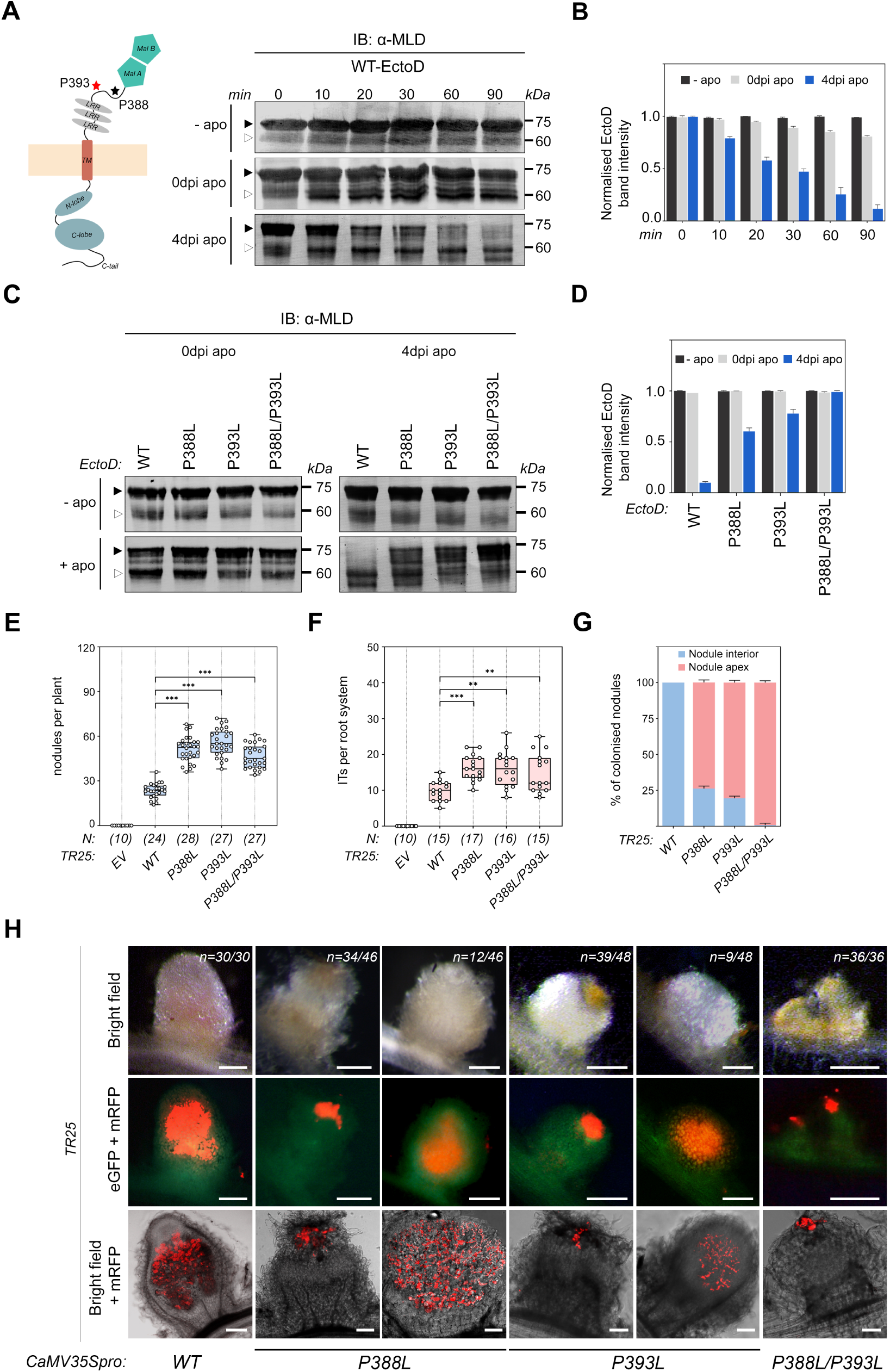
Significance of GDPC+4 Proline in SYMRK EctoD-hinge. (**A-D**) Cleavage of Trx-6xHis tagged SYMRK-EctoD by uninfected (0 dpi) and infected (4 dpi) root apoplastic sap of *Arachis hypogaea*. (**A**) Cleavage at indicated time points for 0 to 90 min was monitored by western blotting with α-MLD. The black and white arrowheads indicate SYMRK EctoD (∼75 kDa) and released MLD (∼60 kDa) respectively, confirmed by LC-MS/MS (see Supplementary Data Set 2). (**B**) Bar plots represents the intensity of ∼75 kDa band quantified using ImageJ. Each bar represents the average intensity of three individual repeats. Error bars represent standard errors of the mean. (**C and D**) Cleavage of Trx-6xHis tagged SYMRK-EctoD mutants (P388L, P393L and P388L/P393L) for 90 min and monitored as described in (A, B). The efficiency of the cleavage being WT> P388L > P393L > P388L/P393L. (**E-H**) Effect of P388L and P393L mutation on nodule development and symbiont colonization. *TR25* complemented with *WT-SYMRK-eGFP*, *P388L-SYMRK-eGFP*, *P393L-SYMRK-eGFP* and *P388L/P393L-SYMRK-eGFP* and infected with *Sinorhizobium meliloti.* IT and Nodule development were monitored in transgenic hairy roots at 1WAI and 6WAI respectively. Box whisker plots represent the total number of nodules/ plant (**E**) and the number of ITs/ root system (**F**). An average of 3 biological replicates each having n > 8 plants for nodule count and n > 4 plants for IT count were analyzed. The number of plants (N) analyzed for each construct are indicated beneath the respective box. Error bar denotes standard deviation (SD). Asterisks denote statistical significance as calculated by using an unpaired two-tailed t test: ***p < 0.001, **p < 0.01, ns, not significant. (**G**) Histogram represents the distribution of rhizobial colonization in nodule apex and nodule interior. (**H**) Stereo images shown as bright-field (upper panels), merged images of eGFP and mRFP (middle panels) and confocal images of merged mRFP and bright field (lower panel). The letter ‘n’ indicates the total number of nodules analyzed for each case. Scale bars, for upper and middle panel, 500μm; for lower panel, 40μm. For details, see Supplementary Fig. S5.

There was a significant phenotypic overlap between *TR25* complemented with KD-hinge (P671E, Y670F) (Fig. 2, D-G) and EctoD-hinge (P388L, P393L) mutants, where in both cases there was significant increase in nodule numbers and IT counts (Fig. 3, E and F). In the majority of the nodules of P388L-SYMRK:*TR25* (n= 34/46) and P393L-SYMRK:*TR25* (n= 39/48), rhizobia were unable to cross the epidermal-cortical barrier and were restricted at the nodule apex (Fig. 3, G and H and Supplementary Fig. S5, D and E, Supplementary Table S1). ITs were ramified in the few remaining nodules, but rhizobial release was never observed. In the double EctoD-hinge mutant P388L/P393L-SYMRK:*TR25* (n=36/36), rhizobial progression was strictly restricted at the epidermal-cortical barrier (Fig. 3, G and H and Supplementary Fig. S5F, Supplementary Table S1). Interestingly, the number of nodules where rhizobia was restricted to the nodule apex in P388L-SYMRK:*TR25* (74%), P393L-SYMRK:*TR25* (81%), and P388L/P393L-SYMRK:*TR25* (100%) shows a clear correlation with the cleavability of the respective EctoD mutants (Figs. 3, C and D vs 3, G and H). The more the EctoD was non-cleavable, the more was the restriction of ITs in the nodule apex indicating cleavage of EctoD to be an essential step in the epidermal-cortical barrier. It may be noted that epidermal IT progression and nodule organogenesis were unaffected by the restriction of EctoD cleavage. This aligns with non-requirement of optimal phosphorylation of SYMRK for initiating the epidermal processes and further affirms the functional involvement of SYMRK at the epidermal-cortical barrier to be distinct from its involvement in the epidermis.

### Double-hinge mutants of SYMRK fail to initiate symbiosis and cause dominant-negative interference at epidermal-cortical barrier

The phenotypic overlap between KD-hinge and EctoD-hinge mutants suggested a crosstalk between hinge-regions of SYMRK. The crosstalk was also evident from observing the restriction of EctoD cleavage in KD-hinge mutants in *Nicotiana* leaves (Supplementary Fig. S6B). To further understand the crosstalk and thereby understand the significance of the adapted Proline residues in SYMRK, we made double-hinge mutants where both the Prolines in KD-hinge (Gatekeeper+1) and EctoD-hinge (GDPC+4) of SYMRK were substituted. Also combination of substitution of other conserved hinge residues were done as indicated. *TR25* complementation with these mutants showed a severe loss of function in all 4 double-hinge mutants (Y670F/P388L-SYMRK:*TR25*, Y670F/P393L-SYMRK:*TR25*, P671E/P388L-SYMRK:*TR25*, and P671E/P393L-SYMRK:*TR25*) with rare signs of ITs and bumps (Fig. 4, A, B and E; Supplementary Table S1). This indicates an intramolecular epistasis between the KD-hinge and EctoD-hinge residues of SYMRK suggesting a hinge-hinge communication in SYMRK to have a role during rhizobial infection. Disruption of the hinge-hinge communication in double mutants appear to affect the conformational dynamics of SYMRK to either block its activation (Yoshida & Parniske, 2005) or prevent its interaction with LysM receptors (Antolín-Llovera, Petutsching, et al., 2014; Krönauer & Radutoiu, 2021) with a complete block in epidermal processes.

**Figure 4.**
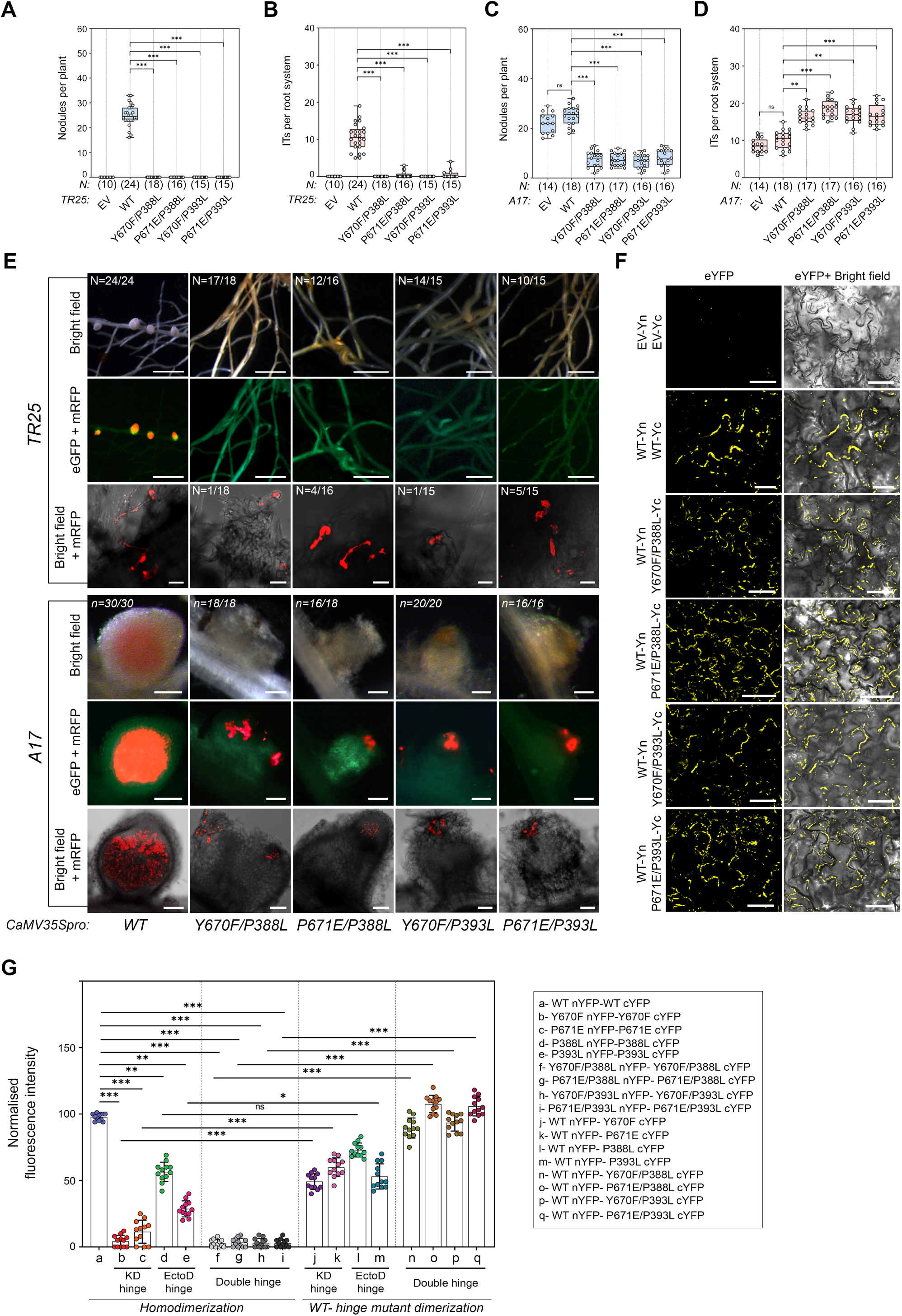
Dominant negative effect of SYMRK double-hinge mutants at the epidermal-cortical barrier. (**A-E**) Effect of double-hinge mutation on nodule development and symbiont colonization. *TR25* (A, B and E) and *A17* (C, D and E) were transformed with double-hinge mutants: Y670F/P388L-SYMRK-eGFP, P671E/P388L-SYMRK-eGFP, Y670F/P393L-SYMRK-eGFP, P671E/P393L-SYMRK-eGFP and infected with *Sinorhizobium meliloti.* IT and Nodule development were monitored in transgenic hairy roots at 1WAI and 6WAI respectively. (**A**-**D**) Box whisker plots represent the total number of nodules/ plant (A, C) and the number of ITs/ root system (B, D). An average of 3 biological replicates each having n > 6 plants for both nodule and IT count were analyzed. The number of plants (N) analyzed for each construct are indicated beneath the respective box. Error bar denotes standard deviation (SD). Asterisks denote statistical significance as calculated using Mann–Whitney U test: ***p < 0.0005 (A, B) and unpaired two-tailed t test: ***p < 0.001, **p < 0.01, ns, not significant (C, D) respectively. (**E**) Stereo images (bright-field and merged eGFP and mRFP) and confocal images (merged mRFP and bright field) in *TR25* (upper 3 panels) indicates rare rhizobial colonization and in A17 (lower 3 panels) indicates dominant negative interference of rhizobial progression at the epidermal-cortical barrier (for details, see Supplementary Fig. S7A). The letter ‘N’ and ‘n’ indicates the total number of plants and total number of nodules analyzed for each case respectively. Scale bars, 2mm (upper and middle panel of *TR25*); 20µm (lower panel of *TR25*) and 500µm (upper and middle panel of *A17*); 40μm (lower panel of *A17*). (**F**, **G**) BiFC assays of WT-SYMRK and SYMRK single hinge and double hinge mutants in indicated combinations for testing their homodimerization or dimerization with WT-SYMRK. (**F**) Confocal images of YFP complementation are shown as eYFP (left panels) and merged images of eYFP and bright field (right panels) for indicated combinations (for additional data, see Supplementary Fig. S8B). Scale bar, 10µm. (**G**) Bar plot represents the normalized YFP fluorescence intensity of KD-hinge, EctoD-hinge and double-hinge mutants of SYMRK for their homodimerization and WT: hinge mutant dimerization. YFP fluorescence intensities from 12 square areas of 150µm x 150µm from 4 different leaf disks of each indicated combinations were quantified using ImageJ software. Fluorescence intensities were normalized with respect to the YFP intensity of *WT-SYMRK-nYFP: WT-SYMRK-cYFP* homodimerization. Error bars denotes standard deviation (SD). Asterisks denote statistical significance as calculated by using one-way ANOVA followed by Tukey’s multiple comparison test: ***p < 0.001, **<0.01, ns, not significant.

To further probe into the mechanism of involvement of the conserved hinge Proline residues in SYMRK symbiotic functions, the double-hinge mutants of SYMRK were introduced in WT *Medicago truncatula* (A17) to check whether they have dominant-negative effect by blocking the normal activity of endogenous SYMRK. Indeed, with all the four KD-hinge:Ecto-hinge double mutants (Y670F/P388L-SYMRK:A17, Y670F/P393L-SYMRK:A17, P671E/P388L-SYMRK:A17, and P671E/P393L-SYMRK:A17) IT progress was hampered at the epidermal-cortical barrier resulting in extensive infection patches on nodule apex (Fig. 4, C, D and E and Supplementary Fig S7, A and C, Supplementary Table S2). Also, in all these cases high IT counts were coupled with low nodule counts indicating a loss of coordination between epidermal and cortical responses. Nodules formed with KD-hinge or EctoD-hinge single mutants resembled the functional red nodules of WT-SYMRK: A17 and the IT or nodule counts were also not affected though there was rare or no release of symbionts (Supplementary Fig S7, B and C, and Supplementary Table S2). The dominant negative effects of double-hinge mutants at the epidermal-cortical barrier but not in the epidermis further reiterates the distinct nature of involvement of SYMRK in epidermis and epidermal cortical barrier. These difference can originate from the difference in multimeric receptor complexes formed in each barriers (Krönauer & Radutoiu, 2021).

Earlier we have noted dimerization of SYMRK that was affected by gatekeeper substitutions within the KD-hinge (Paul et al., 2014). We therefore envisaged the dominant-negative interference to be due to ineffective hybrid dimer formation between the double-hinge mutants and the endogenous SYMRK. To test this hypothesis, we implemented bimolecular fluorescence complementation (BiFC) in *Nicotiana* leaves. Reconstitution of YFP fluorescence clearly indicated dimerization of WT-SYMRK (Fig. 4F). It may be noted that though SYMRK as well as its single and double-hinge mutants homogeneously localised in plasma membrane (PM) (Supplementary Fig. S8A), the YFP fluorescence from BiFC was distributed into patches of varying sizes ranging from small punctae to extensive patches (Fig 4F). There are reports of microenvironments supporting assembly of receptors where membrane lipids are capable of molecular interactions with the transmembrane domains of receptors, or the receptor binds to proteins with intrinsic dimerization potential to guide the assembly (Li et al., 2024). Irrespective of the underlying mechanism, reconstituted YFP clearly indicated WT-SYMRK to homodimerize in the absence of familiar settings of rhizobia-legume interaction, most likely in a ligand-independent manner.

Next we checked the dimerization of KD-hinge, EctoD-hinge, and double-hinge mutants of SYMRK (Fig 4F, and Supplementary Fig. S8B). Both homodimerization and their dimerization with WT-SYMRK differed significantly as reflected in a quantitative readout of the dimerization potential of all hinge mutants in terms of the frequency and intensity of fluorescence in cells expressing YFP (Fig 4, F and G, and Supplementary Fig. S8B). The readout was consistent and emerged with a pattern. Homodimerization was not detectable in any of the double-hinge mutants but their efficiency of hybrid dimer formation with WT-SYMRK was significantly higher than KD-hinge or EctoD-hinge mutants. Formation of these nonfunctional hybrid dimers of SYMRK with double-hinge mutants gives a potential explanation for the observed dominant-negative interference at the epidermal-cortical barrier in A17. Since double mutants did not interfere in the initiation of ITs and nodule organogenesis, the participation of SYMRK in the epidermis appeared to be monomeric within heteromeric complexes where endogenous receptor could generate successful signal. This affirms the widely accepted model of monomeric SYMRK participation in the epidermis (Antolín-Llovera, Petutsching, et al., 2014; Krönauer & Radutoiu, 2021). On the other hand, dimerization of SYMRK appear necessary at the transcellular epidermal-cortical barrier. This aligns with the requirement of optimal phosphorylation of SYMRK at the same barrier (Fig. 2) as receptor kinases exploit dimerization and/or multimerization for direct regulation of their kinase domains (Lavoie et al., 2014).

The readout of dimerization of single-hinge mutants of SYMRK allow important derivations related to the major events at the epidermal cortical barrier. Significant homodimerization in EctoD-hinge mutants indicate cleavage is not necessary for dimerization whereas insignificant homodimerization in KD-hinge mutants indicated kinase domain to be the main architect of dimerization (Fig. 4G and Supplementary Fig. S8B). This aligns with the current view that closed conformation of active KDs offer stable interface propitious to dimerization (Lavoie et al., 2014). Since KD-hinge mutants restricted cleavage of SYMRK EctoD (Supplementary Fig. S6B), it is possible that absence of dimerization in KD-hinge mutants have restricted optimal activation of SYMRK for activating EctoD cleavage. The interference in the endocytic barrier by the single-hinge mutants of SYMRK in A17 (Supplementary Fig. S7) is not understood but was most likely due to negative interference of inappropriate phosphocode or unshedded EctoD in the multimeric receptor complexes.

### Released MLD has an active role in symbiosis: Restriction in progress of symbionts by hinge mutants of SYMRK is overcome in presence of MLD

The key molecular events associated with SYMRK in the epidermal-cortical barrier is its dimerization, optimal phosphorylation and EctoD cleavage. If these events constitute a molecular module for guarding the transcellular progress of symbionts across this barrier, the question was whether the primary objective of evolving this functional module was to release MLD as an active principle for guiding symbiosis. If so, we reasoned that co-expressing free MLD should be able to bypass the restriction caused by the hinge mutants of SYMRK. For this we used a binary destination vector for simultaneous ectopic expression of MLD (1 to 350 aa) along with the KD-hinge or EctoD-hinge and the double-hinge mutants of SYMRK in *TR25* (Supplementary Fig. S9, A and B). In presence of MLD, there was a significant increase in nodule numbers in *TR25* complemented with KD-hinge and EctoD-hinge mutants (Fig 5A and Supplementary Table S1). Almost all nodules were pink similar to those that developed in MLD/WT-SYMRK:*TR25* without any infection patches on the nodule apex (Fig 5, B and C and Supplementary Fig. S9C). Even the double EctoD-hinge mutant P388L/P393L-SYMRK that was completely resistant to cleavage and strictly restricted symbionts at the epidermal-cortical barrier demonstrated complete crossover in presence of MLD. This complete crossover of epidermal-cortical boundary by the symbionts in both KD-hinge and EctoD-hinge mutants directly indicated a role of released MLD in symbiosis signalling and indicated the modular nature of SYMRK action. Crossover in EctoD-hinge mutants in presence of MLD indicated uncleaved SYMRK was not a hindrance in the epidermal-cortical barrier. Whereas, crossover in KD-hinge mutants in presence of MLD indicated the phosphocodes generated in WT-SYMRK to be primarily required for its release. Therefore, KD-hinge mediated optimal activation and EctoD-hinge mediated EctoD cleavage of SYMRK was necessary but not essential at the epidermal-cortical barrier if free MLD was available. Unexpectedly, the apparently functional pink nodules developed in presence of MLD showed rare release of symbionts (Supplementary Fig. S9C) suggesting both components of SYMRK signalling is necessary and essential in the endocytic barrier as free MLD was barely able to restore the crossing of endocytic barrier.

**Figure 5.**
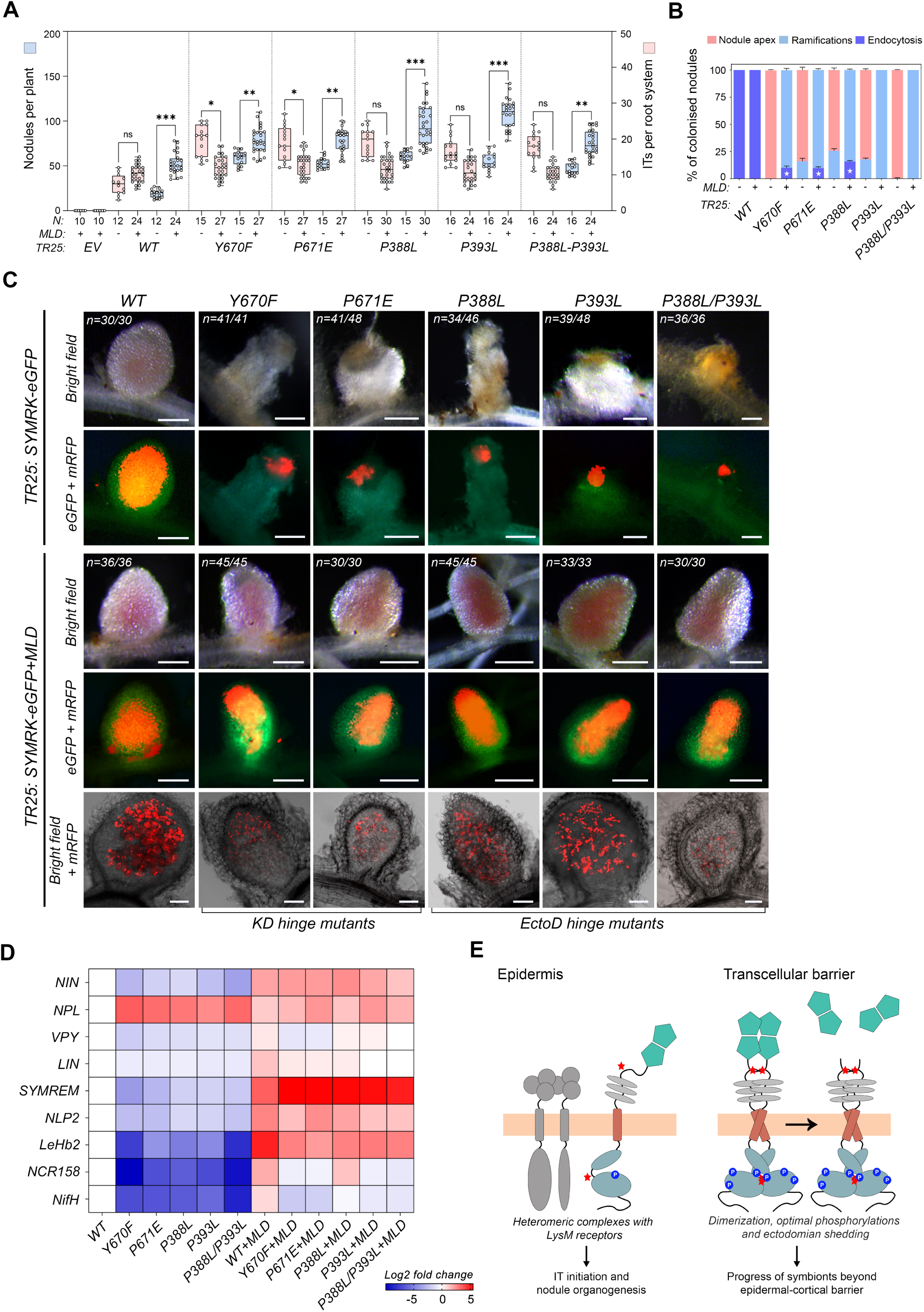
Expression of MLD overcomes the restriction of rhizobial progress at the epidermal-cortical barrier caused by SYMRK hinge mutants. (**A-C**) Effect of MLD coexpression on nodule development and symbiont colonization in *TR25* complemented with hinge mutants of SYMRK. *TR25* complemented with binary expression vector containing *ROLDpro:WT-SYMRK-eGFP* or *ROLDpro:SYMRK-hinge mutants-eGFP* with or without simultaneous expression of *CaMV35Spro:MLD* and infected with *Sinorhizobium meliloti.* IT and Nodule development were monitored in transgenic hairy roots at 1WAI and 6WAI respectively. (**A**) Box whisker plots represent the total number of nodules/ plant (left Y axis) and the number of ITs/ root system (right Y axis). An average of 3 biological replicates each having n > 4 plants for both nodule and IT count were analyzed. The number of plants (N) analyzed for each construct are indicated beneath the respective box. Error bar denotes standard deviation (SD). Asterisks denote statistical significance as calculated by using an unpaired two-tailed t test: ***p < 0.001, **p < 0.01, ns, not significant. (**B**) Histogram represents the distribution of rhizobial colonization in nodule interior. (**C**) Stereo images (bright-field and merged eGFP and mRFP) and confocal images (merged mRFP and bright field) (for details, see Supplementary Fig. S9C) indicate ramifications of ITs in the nodule interior in presence of MLD, which was otherwise restricted at the epidermal-cortical barrier in absence of MLD, forming infection patches at nodule apex. The letter ‘n’ indicates the total number of nodules analyzed for each case. Scale bars, 500µm (for stereo images); 40µm (for confocal images). (**D**) RT-qPCR analysis of indicated gene in nodulated roots of *TR25* at 2WAI, transformed with binary expression vector containing *ROLDpro:WT-SYMRK-eGFP* or *ROLDpro:SYMRK-hinge mutants-eGFP* with or without simultaneous expression of *CaMV35Spro:MLD*. Heatmap represents an average of 3 biological replicates each having n > 5 plants. Fold changes are indicated in log_2_ scale, p, <0.05 (for individual bar plots, see Supplementary Fig. S10). (**E**) Working model of SYMRK action at epidermal-cortical barrier. At epidermis, SYMRK functions in monomeric form within heteromeric complexes which is a widely accepted model for SYMRK action. The key molecular events associated with SYMRK in the epidermal-cortical barrier are its (i) dimerization, (ii) optimal phosphorylation and (iii) EctoD cleavage. Conserved Proline residues in SYMRK orthologues in gatekeeper +1 and GDPC+4 positions (indicated by ‘red’ stars) have critical roles in each of these molecular events. None of these events were important in the epidermis for initiation of infection threads or nodule organogenesis indicating their importance in cross over of transcellular barriers.

To check how far the symbiosis was restored in *TR25* complemented with SYMRK hinge mutants in presence of MLD, we checked the expression of markers that have a tight connection with N_2_ fixation like *NODULE INCEPTION (NIN), NIN-LIKE PROTEIN (NLP2), LEGHAEMOGLOBIN (LeHb2), NODULE CYSTEINE-RICH PEPTIDE (NCR158) and NITROGENASE REDUCTASE (NifH).* NIN-like protein NLP2 and NIN directly activate the expression of LeHb2 for maintaining micro-oxic environment to protect Nitrogenase, and NCR158 is known to express in the nitrogen fixing zone (Jiang et al., 2021). Expression of *MtNIN* and *MtNLP2* was significantly low in *TR25* complemented with hinge mutants whereas expression of *MtLeHb2*, MtNCR*158* and *NifH* were undetectable (Fig. 5D and Supplementary Fig.10). In presence of MLD, expression of *MtNIN* and *MtNLP2* were significantly higher than the corresponding hinge mutants complemented *TR25* and accordingly the expression of *MtLeHb2, MtNCR158* and *NifH* was also restored to almost normal level. Restoration of these indicators in the hinge mutants of SYMRK in presence of MLD indicates restoration of symbiosis by the infrequent release of symbionts that was observed. But considering the rarity of release it remains to be seen whether observed phenotype is close to fixation thread symbiosis. We also checked the expression of markers associated with IT initiation and propagation like NODULE PECTATE LYASE (NPL), VAPYRIN (VPY), LUMPY INFECTIONS (LIN), and SYMBIOSIS ASSOCIATED REMORIN (SYMREM1). IT counts were significantly higher in all hinge mutants and their progress was hindered with inflated sacs but in presence of MLD the number of ITs matched the counts in WT-SYMRK:*TR25* and their progress was normal (Fig 5A and Supplementary Table S1). NPL regulates degradation of de-esterified pectins and thereby supports IT initiation and propagation (Su, Zhang, et al., 2023). Accordingly, *MtNPL* expression was relatively higher (∼6-12 fold) in absence of MLD in all hinge mutants and remained high in presence of MLD (∼4-6 fold) because of extensive ramification of ITs in the nodule interior (Fig. 5D and Supplementary Fig.10). VPY and LIN maintains the polar growth of ITs (Lace et al., 2023) and their significantly low expression in *TR25* complemented with hinge mutants of SYMRK explain the anomalous progress of ITs. Expression of both *MtVPY* and *MtLIN* was restored in presence of MLD (Fig. 5D and Supplementary Fig.10). SYMREM1 a symbiosis specific remorin, has a role in recruiting and stabilising the symbiotic receptors in membrane nanodomains near cell wall depleted tips of ITs that is essential for its transcellular migration (Su, Rodriguez-Franco, et al., 2023). In presence of MLD, *MtSYMREM1* expression was significantly higher than in both KD-hinge and EctoD-hinge mutants complemented *TR25* (∼200-500 fold) and therefore could be crucial for the crossover of ITs in hinge mutants of SYMRK. It may be noted that the rare formation of ITs in *TR25* complemented with double-hinge mutants of SYMRK was not affected in presence of MLD which is in accordance with no effect of MLD on symbiotic gene expression (Supplementary Fig. S10).

In presence of MLD, extracellular matrix mechanical cues can be envisaged to generate a niche for transcellular infection threads migration but how apoplastic release of MLD functions to reset the symbiosis associated gene expression is not understood at this point. But the remarkable effect of released MLD in the progress of symbiosis in the hinge mutants of SYMRK, strongly suggests the KD-hinge and EctoD-hinge regions together to have evolved a module for release of MLD.

## Discussion

Taken together, our observations highlight the functional significance of two distinct Proline residues present in the hinge-region of SYMRK orthologues which is in accordance with hinge motions playing a significant role in the functional evolution of proteins (Campitelli et al., 2020). Both Prolines qualifies for “what must be acquired or cannot be lost for a functional symbiosis” (Zhang et al., 2024). Among all MLD-LRR-RLKs only SYMRK orthologues have a distinct Proline in gatekeeper+1 position of KD-hinge, which is important for dimerization and generation of an optimal phosphocode (Figs. 1, 2 and 4). Adaptation of this Proline appears crucial for evolving and defining SYMRK. This ‘Gatekeeper+1Pro’ event in SYMRK probably triggered assembly of CSSP in land plants because its pro-orthologue MLD-LRR-RLK in Zygnematophyceae has a Glutamine in the same position. Since MLD-LRR-RLKs are involved in immune responses (Yeh et al., 2016) this event in SYMRK could have allowed neo-functionalization specific to symbiotic responses. Subsequently within NFC, there was adaptation of another distinct Proline in ectoD-hinge region of SYMRK orthologues at GDPC+4 position within a conserved W^1^xGDPC^1^x_3_PW^2^xGx_2_C^2^ motif. Associated with this gain of Proline at GDPC+4, there was a conspicuous loss of Proline in GDPC+2 position thereby giving a distinct signature ‘GDPC+2Pro^-^/4Pro^+^’ for ectoD-hinge in SYMRK orthologues within legumes that determined the cleavability of ectodomain within the motif (Figs. 1 and 3). However, apart from determining the cleavability of ectodomain this motif might function as an integrin like thiol switch (Lorenzen et al., 2021) responding to environmental cues as indicated by the predicted open and closed structure of the motif and its effect on ectodomain structure of SYMRK (Supplementary Fig. S3). Whatever be the underlying mechanism, since RNS has evolved as a stable trait in legumes this ‘GDPC+2Pro^-^/4Pro^+^’ event appear to be a legume specific adaptation. The KD-hinge:EctoD-hinge crosstalk in SYMRK act as a signalling module for releasing MLD, where dimerization and optimal phosphorylation of SYMRK determines ectoD cleavage (Figs. 4 and 5). The released MLD allow cross over of epidermal-cortical barrier by the symbionts upon activating symbiosis related gene expression (Fig. 5).

Based on our findings and what is known in the field we propose SYMRK signalling to differ in the epidermis and in the epidermal-cortical barrier (Fig. 5E). In the epidermis, ITs are invaginations of the cell wall and plasma membrane whereas in the epidermal-cortical barrier or in the endocytic barrier, ITs cross transcellular barriers. Signalling for epidermal symbiotic responses can originate from a monomeric intact SYMRK as disruption of cleavage did not affect the epidermal processes of initiation and nodule organogenesis (Fig. 3). This aligns and affirms the established model of SYMRK signalling where it is a coreceptor with LysM RLKs that senses the lipo chitooligosaccharides secreted by its nitrogen-fixing microsymbiont and activates the CSSP (Antolín-Llovera, Petutsching, et al., 2014; Krönauer & Radutoiu, 2021). Such heteromeric receptor complexes with LysM receptors is also demonstrated with MLD-LRR-RLKs (Yeh et al., 2016). On the other hand, cleavage of EctoD and release of MLD in response to dimerization and optimal phosphorylation, is essential at the epidermal-cortical barrier. Both ‘Gatekeeper+1Pro’ event in KD-hinge and the ‘GDPC+2Pro^-^/4Pro^+^’ event in ectoD-hinge are crucial for each of these key events. It’s an open question how dimerization of SYMRK was restrained in the epidermis or how it was facilitated at the transcellular barrier. Finally, it was intriguing to note that the Proline signature in ectoD-hinge region of ceasalpinoids and mimosoids resembled the signature in non-legumes within NFC instead of their legume partners. In the backdrop of widely acclaimed role of SYMRK in endocytic accommodation of symbionts it remains to be seen whether the exclusive legume specific innovation in SYMRK ectodomain has a role in intracellularisation of symbionts within symbiosomes.

## Materials and Methods

### Multiple sequence alignment and phylogenetic tree analysis

Protein sequences of LRR-RLK-I, MLD-RLKs and LRR-RLK-II members were retrieved from the National Center for Biotechnology Information (NCBI) database (https://www.ncbi.nlm.nih.gov/), UniProt (https://www.uniprot.org/) or by BLASTp search against *Ah*SYMRK. SYMRK orthologous sequences were obtained from (Radhakrishnan et al., 2020). Multiple sequence alignment and maximum likelihood phylogenetic trees (using Jones-Taylor-Thontin model, 1000 bootstrap replicates) of full length, kinase domain and Ectodomain was developed using MEGA-X software. Visualization and color-code annotation of the trees were accomplished using Interactive Tree of Life (iTOL).

### Structural analysis

SYMRK structural models were obtained from AlphaFold2 online server (Jumper et al., 2021). Predicted structural models of SYMRK from *Marchantia paleacea* (KAG6549668.1), LRR-RLK-I from *Marchantia polymorpha* (PTQ39574.1), SYMRKL1 of *Lotus japonicus* (LotjaGi2g1v0191100) were generated in ColabFold v1.5.5: AlphaFold2 using MMseqs2. ColabFold66 (Mirdita et al., 2022). Default settings were utilized (use_amber: no, template_mode: none, msa_mode: MMSeqs2, pair_mode: unpaired+paired, model_type: auto, num_recycles: 3). Structural alignments were done in PyMOL (The PyMOL molecular graphics system, version 3, Schrödinger, LLC).

### Plant materials and Growth conditions

*Medicago truncatula TR25* and *A17* seeds were grown and infected with Sinorhizobium *meliloti Sm2011-pBHR-mRFP* as described in (Molla et al., 2023). *Arachis hypogaea* JL24 seeds were grown and infected with Semia-6144 as described in (Sinharoy & DasGupta, 2009). In both cases germinated seedlings were placed in Fahraeus 0.8% agar (w/v) medium in square petri dishes and maintained at 20°C for *Medicago* and 25°C for *Arachis* on a 16h-8h light-dark schedule under 100 to 110 μmol/m^2^/s for further growth. *Nicotiana benthamiana* seeds were grown in 1:1 Soilrite:Vermiculite at 25°C on a 16h-8h light-dark schedule under 200 μmol/m^2^/s.

### Plasmid constructs

All constructs are listed in Supplementary Table S4. *Arachis hypogaea* SYMRK construct (*CaMV35Spro:WTSYMRK-eGFP-t35S*) was used as template for generating *AhSYMRK* constructs as described before (Saha et al., 2016). Primers are listed in Supplementary Table S3. For generating vectors containing two expression cassettes PCR fragments with flanking attB sites were first recombined with pDONR221-P3/P4 to obtain pENTR L3-L4 clones using Gateway™ BP Clonase™ II Enzyme mix (ThermoFisher Scientific) which was then recombined in the double vector pK7m34GW2-8m21GW3 (Karimi et al., 2007) using Gateway™ LR Clonase™ II Enzyme mix (ThermoFisher Scientific) to create *ROLDpro:SYMRK-eGFP-tOCS, CaMV35Spro:signalpeptide+MLD-t35S* constructs. For recombinant protein production, SYMRK kinase domain and SYMRK ectodomain were cloned into pET28a and pET32a to generate 6xHis tagged SYMRK-KD and Trx-6xHis tagged SYMRK-EctoD respectively. For BiFC analysis, all entry clones of SYMRK were recombined into two destination vectors, pSAT4(A)-DEST-nEYFP-N1(pE3134) and pSAT5(A)-DEST-nEYFP-N1(pE3132) using Gateway™ LR Clonase™ II Enzyme mix (ThermoFisher Scientific) to create a) *CaMV35Spro: SYMRK-n-term (1-174) EYFP-t35S* and b) *CaMV35Spro: SYMRK-c-term (175-251) EYFP-t35S* constructs.

### Recombinant protein purification and analysis

Recombinant SYMRK-KD purification was done as described before (Bhattacharya et al., 2019; Samaddar et al., 2013). Recombinant SYMRK-EctoD proteins were purified from *E. coli* inclusion body pellet. Cells were incubated overnight at 4°C in lysis buffer containing 50mM Tris-Cl, pH 8.0; 5mM EDTA; 5mM βME; 5mM PMSF; 1mg/ml of lysozyme before sonication and centrifugation at 10000×g for 30 min. The pellet was washed 5-6 times with wash buffer containing 50mM Tris-Cl, pH 8.0; 300mM NaCl; 5mM βME, 2M urea, 2% Triton-X-100 and were then resuspended overnight in extraction buffer containing 50mM Tris-Cl, pH 8.0; 300mM NaCl; 5mM imidazole; 5mM βME and 8M urea. Undissolved particles were removed by centrifugation and supernatant was loaded on to Ni-NTA column. Column was washed with equilibration buffer containing 50mM Tris-Cl, pH 8.0; 300mM NaCl; 20mM imidazole; 5mM βME and 8M urea and proteins were eluted using elution buffer containing 50mM Tris-Cl, pH 8.0; 300mM NaCl; 500mM imidazole; 5mM 2-mercaptoethanol and 8M urea. Eluted proteins were dialyzed in a buffer containing 50mM sodium phosphate buffer, pH 6.0; 10% glycerol. Protein concentration was estimated by using Quick Start™ Bradford Protein Assay (Bio Rad). 2D gel electrophoresis was performed as described previously (Bhattacharya et al., 2019) using 11 cm Immobiline DryStrips pH 4-7 (GE Healthcare) in Ettan IPGphor3 IEF system (GEHealthcare). Western blot was performed as described previously (Bhattacharya et al., 2019). Membranes were probed with α-pSer (Sigma, 1:1000 dilution); α-pTyr (Cell Signaling Technology, 1:3000 dilution); α-pThr (Cell Signaling Technology, 1:3000 dilution), and two custom-made polyclonal antibodies α-pY670 (Imgenex India, 1:5000 dilution) and α-MLD (Biobharati lifescience, 1:5000 dilution).

### Sample preparation and LC-MS/MS analysis

For phosphopeptide analysis, purified 6xHis-SYMRK-KD proteins were *in vitro* phosphorylated and subjected to in-sol trypsin (Promega) digestion. Phosphopeptide enrichment was done as described before (Saha et al., 2016). SYMRK-KD phosphopeptides were analyzed in Xevo G2-XS QTof Quadrupole Time-of-Flight Mass Spectrometer (Waters). Injected peptides were separated and eluted by Acquity UPLC Beh C18 Column (130Å, 1.7 µm, 2.1 mm X 50 mm, Waters) with 55 min linear gradient of 5 to 45% mobile phase B (0.1% FA in 95% MeCN) against mobile phase A (0.1% FA in 95% H_2_O) at a flow rate 50ul/min. Instrument was set to positive ionization mode. Precursor ion (MS) acquisition range was adjusted to 100–2000 m/z at 5 spectra/sec with capillary voltage 2.5kV and cone voltage 70V. The most intense precursor ions were selected for fragmentation with higher-energy collisional induced dissociation (CID, 18-40 V). The raw data files were processed using Progenesis QI (Waters) software. Peak lists were searched against *Arachis hypogaea* UniProt database (https://www.uniprot.org/) appended with the SymRK wild type and mutant sequences. The following parameters were selected to identify tryptic peptides for protein identification: 10 ppm precursor ion mass tolerance; 0.6 Da product ion mass tolerance for CID and a maximum of two missed cleavages; carbamidomethylation of Cys was set as a fixed modification; oxidation of Met and phosphorylation of Ser, Thr, and Tyr were set as variable modifications. MS/MS spectra for phosphopeptides were generated using ProteinLynx Global Server (PLGS).

### Ectodomain cleavage assay

Uninfected (0dpi) and infected with *Bradyrhizobium sp. SEMIA 6144* (4dpi) *Arachis hypogaea* roots were used for preparation of apoplastic extracts as described in (Zhou et al., 2011). In brief, roots were vacuum infiltrated for 15 min in buffer containing 100mM Tris-HCL, pH 7.5; 200mM KCl; 1mM PMSF. Roots were then cut at the root-shoot junction and centrifuged for 15 min at 1500g at 4°C to collect the apoplastic sap. The cytoplasmic contamination of the sap was calculated as the percentage of G6PDH activity in the apoplast as compared with the activity in the total protein extract, which was < 5%. Extracted apoplastic sap were used as a source of proteases for *in vitro* Ectodomain-cleavage assay. For this, purified SYMRK EctoDomains were incubated with apoplastic sap in a reaction buffer (50 mM sodium phoshphate buffer, 1 mM EDTA, pH 6.0) at 37°C and stopped with 1x Laemmli buffer. The reaction mixture was subjected to western blot with custom made anti-MLD antibody (α-MLD: Biobharati lifescience, 1:5000 dilution). Both the cleaved and un-cleaved bands of SYMRK EctoD were then subjected to in gel tryptic digestion and peptide extraction as described in (Saha et al., 2016). Peptides were analysed in LTQ Orbitrap XL mass spectrometer (ThermoFisher Scientific).

### Generation of composite *M. truncatula* plants and phenotypic analysis

*Agrobacterium rhizogenes* mediated hairy root transformation was done based on the protocol described previously (Molla et al., 2023; Saha et al., 2016). Leica stereo fluorescence microscope M205FA equipped with a Leica DFC310FX digital camera (Leica Microsystems) was used to acquire images of whole-mount nodulated roots. Confocal Microscopy was done using a TCS SP5 II AOBS confocal laser scanning microscope (Leica Microsystems). All digital micrographs were processed using Adobe Photoshop 2020. RNA extraction, cDNA synthesis, and quantitative reverse-transcription polymerase chain reaction were done as described in (Molla et al., 2023).

### Transient protein Expression and BiFC analysis in *Nicotiana benthamiana*

*Agrobacterium tumefaciens* GV3101 strains were transformed with the vectors carrying *CaMV35Spro:SYMRK-eGFP* (for localization) and *CaMV35Spro:SYMRK-nYFP + CaMV35Spro:SYMRK-cYFP* (for BiFC) grown on YEP at 28°C. For agro-infiltration, cell pellet was resuspended in buffer containing 10mM MgCl_2_, 10mM MES, 150µM acetosyringone pH 5.6, to an OD_600_=0.5. GV3101 strains expressing a p19 RNA silencing suppressor (OD_600_=0.5) were mixed with GV3101 strains containing SYMRK constructs, 1:1 for localization and 1:1:1 for BiFC Split-YFP assays. In both cases mixtures were syringe-infiltrated in the leaves from 4 to 5-week old *N. benthamiana* plants. After 70-72h infiltration, leaf disks were cut and placed on a glass slide with abaxial surface facing upwards and covered by a coverslip. eGFP fluorescence (excitation/emission 488/507 nm) or YFP fluorescence (excitation/emission 513/527 nm) was captured with Leica TCS SP5 II AOBS confocal laser scanning microscope (Leica Microsystems). For BiFC, YFP fluorescence intensities from 12 square areas of 150µm x 150µm from 4 different leaf disks of each experiments were quantified using ImageJ software. Fluorescence intensities were normalised with respect to the YFP intensity of *WT-SYMRK-nYFP:WT-SYMRK-cYFP* homodimerization.

For protein analysis, 5-6 leaf disks expressing SYMRK constructs were ground in liquid nitrogen and resuspended in 500µl homogenization buffer [50mM HEPES, pH 7.5; 150mM NaCl; 5mM EDTA; 5mM EGTA; 10% glycerol; 5mM DTT; 10mM NaF; 10mM Na_3_VO_4_; 1mM PMSF; 1% PVP; protease inhibitor cocktail (sigma)] and kept in ice for 45min. Then centrifuged at 5000g for 10 min at 4°C to remove the debris. The supernatant was ultra-centrifuged at 1,35,000g for 60min at 4°C. The pellet (microsomal fraction) was resupended in 100ul of membrane solubilizing buffer [25mM HEPES, pH 7.5; 100mM NaCl; 5mM EDTA; 5mM EGTA; 10% glycerol; 1% Triton-X-100; 2.5mM DTT; 5mM NaF; 5mM Na_3_VO_4_; 1mM PMSF; 0.5% PVP; protease inhibitor cocktail (sigma)]. Samples were then boiled with 1x laemmli buffer and western blotted with custom made anti-SYMRK_KD_ antibody (Imgenex India, 1:5000 dilution).

### Statistical analysis

All the statistical analysis was carried out using the GraphPad Prism 8.0.2 software. Box limits in the graphs represent 25th to 75th percentile, the horizontal line is the median. Every graph displays all data points. The exact statistical test used for each dataset is indicated in the corresponding figure legend. Statistical analyses were performed as described in each figure legend. Statistical data are provided in Supplementary Data Set 3.

### Accession numbers

Sequence data from this article can be found in the GenBank/EMBL data libraries under accession numbers *Mt*NIN, Medtr5g099060; *MtNPL*, Medtr3g086320; *MtVPY*, Medtr6g027840; *MtLIN*, Medtr1g090320; *MtSYMREM,* JQ061257; *MtNLP2,* Medtr4g068000*; MtLeHb2,* Medtr5g081030; *MtNCR158,* Medtr7g027180; *SmNifH,* AHK23441.

## Acknowledgments

We thank Pierre-Marc Delaux for providing sequences of SYMRK orthologues and Jan de Vries and Yanbin Yin for providing the sequence of pro-orthologue of SYMRK in Zygnematophyceae. We also thank Giles Oldroyd and Christian Rogers for *TR25* seeds, Ton Bisseling and Erik Limpens for *S. meliloti* harboring pBHR-mRFP, Douglas R. Cook for Agrobacterium strain MSU440. We thank Dr. Senjuti Sinharoy for critical discussions; Dona Ghosh, Sampurna Chakraborty and Soumyadeep Saha for their support in cloning and maintaining transgenic plants. We also thank Souvik Roy of Bose Institute, Kolkata, India and Santu Paul of IICB, Kolkata, India for their technical assistance with the LC-MS/MS experiments. For funding, we thank DBT-CEIB (BT/01/CEIB/09/VI/10), DST (EMR/2015/001006), DST-SERB (JCB/2019/000003), CEFIPRA (IFC-6303-2).

## Author Contributions

M.D.G. conceived the project. M.D.G. and D.C. designed experiments and wrote the manuscript, which all other authors read and provided feedback. D.C. performed and coordinated most of the experiments and together with A.P. did the biochemical experiments. A.P. together with S.B. and D.C. demonstrated the dominant negative phenotypes. F.M. contributed in generating and analysing transgenic plants. A.P., S.B., and F.M. assisted D.C. with confocal microscopy. D.C. and S.D. performed the *Nicotiana* infilatration and BiFC assays. A.B. contributed to apoplastic sap extraction.

## Supplementary Data

Supplementary Figure S1. Sequence similarity trees (Supports Fig.1).

Supplementary Figure S2. Alignment of KD-hinge sequences (Supports Fig.1).

Supplementary Figure S3. Hinge region of *Ah*SYMRK ectodomain (Supports Fig.1).

Supplementary Figure S4. Alignment of EctoD-hinge sequences (Supports Fig.1).

Supplementary Figure S5. Detailed phenotype of TR25 complemented with KD-hinge mutants (Supports Fig.2 and 3).

Supplementary Figure S6. SYMRK ectodomain cleavage (Supports Fig.3).

Supplementary Figure S7. Detailed phenotype of A17 transformed with hinge mutants (Supports Fig.4).

Supplementary Figure S8. BiFC assays demonstrating dimerization of SYMRK (Supports Fig.4).

Supplementary Figure S9. Effect of ectopic expression of MLD in TR25 complemented with SYMRK hinge mutants (Supports Fig.5).

Supplementary Figure S10. RT-qPCR analysis of symbiotic genes (Supports Fig.5).

Supplementary Table S1. Infection and nodulation counts in TR25 roots transformed with SYMRK constructs.

Supplementary Table S2. Infection and nodulation counts in A17 roots transformed with SYMRK constructs.

Supplementary Table S3. List of primers used for cloning and RT-qPCR.

Supplementary Table S4. List of constructs used in this study.

Supplementary Data Set 1. List of phosphopeptides of SYMRK-KD and its KD hinge mutants quantitated by LC-MS/MS (Q-TOF).

Supplementary Data Set 2. List of identified peptides of SYMRK-EctoD bands from in vitro cleavage assay quantitated by LC-MS/MS (Q-TOF and Orbitrap).

Supplementary Data Set 3. Results of statistical analysis.

